# ABA signaling prevents phosphodegradation of the *Arabidopsis* SR45 splicing factor to negatively autoregulate inhibition of early seedling development

**DOI:** 10.1101/2022.02.03.479016

**Authors:** Rui Albuquerque-Martins, Dóra Szakonyi, James Rowe, Alexander M. Jones, Paula Duque

**Affiliations:** Instituto Gulbenkian de Ciência, 2780-156 Oeiras, Portugal; Sainsbury Laboratory, University of Cambridge, Cambridge B2 1LR, UK

**Author notes:** **Correspondence:** Paula Duque; Alexander M. Jones.

**Keywords:** Abscisic acid (ABA), alternative splicing, *Arabidopsis thaliana*, protein phosphorylation, SR proteins

## Abstract

Alternative splicing is a key posttranscriptional mechanism to expand the coding capacity of eukaryotic genomes. Although the functional relevance of this process remains poorly understood in plant systems, major modulators of alternative splicing called serine/arginine-rich (SR) proteins have been implicated in plant stress responses mediated by the abscisic acid (ABA) hormone. Loss of function of the *Arabidopsis thaliana* SR-like protein SR45, a *bona fide* splicing factor, has been shown to cause plant hypersensitivity to environmental cues and activation of the ABA pathway. Also, consistent with both animal and plant SR proteins being extensively and reversibly phosphorylated at their C-termini, ABA-induced changes in the phosphorylation status of SR45 have been reported.

Here, we show that *SR45* overexpression reduces *Arabidopsis* sensitivity to ABA during early seedling development. Moreover, exposure to ABA dephosphorylates SR45 at multiple amino acid residues and leads to accumulation of the protein via reduction of SR45 ubiquitination and proteasomal degradation. Using phosphomutant and phosphomimetic transgenic *Arabidopsis* lines, we demonstrate the functional relevance of ABA-mediated dephosphorylation of a single SR45 residue, T264, in antagonizing SR45 ubiquitination and degradation to promote its function as a repressor of seedling ABA sensitivity. Taken together, our results reveal a mechanism in which ABA signaling negatively autoregulates during early plant development via posttranslational control of the SR45 splicing factor.

## INTRODUCTION

Serine/arginine-rich (SR) proteins are members of a highly conserved family of RNA-binding factors that play important roles in mRNA splicing. These accessory spliceosomal proteins bind to *cis-*regulatory sequences in the precursor-mRNA (pre-mRNA), promoting early spliceosome assembly and influencing splice site selection. SR proteins are thus crucial in alternative splicing, a posttranscriptional mechanism that generates multiple transcripts from the same gene and has key biological relevance in eukaryotes, namely in developmental programs and stress response. Structurally, SR proteins present one or two N-terminal RNA recognition motifs (RRMs), responsible for binding transcripts, and an arginine/serine-rich (RS) protein-protein interaction domain at their C terminus that recruits core spliceosome components to splice sites. The RS domain is subjected to extensive reversible phosphorylation (reviewed in Duque, 2011), which in animal systems is known to be required for spliceosome assembly and has also been linked to mRNA export from the nucleus (Huang et al., 2004; Sanford et al., 2005) or in switching functions from splicing activator to repressor (Shi and Manley, 2007; Feng et al., 2008). In plant systems, SR proteins are being increasingly implicated in abiotic stress responses mediated by the abscisic acid (ABA) hormone (Carvalho et al., 2010; Chen et al., 2013; Xing et al., 2015; Albaqami et al., 2019; Laloum et al., 2021). The phytohormone ABA coordinates several developmental processes, such as seed dormancy, embryo maturation, seedling growth and greening, but is also pivotal in the response to environmental challenges. Sensing of osmotic or oxidative stress by plant cells leads to a massive increase in the levels of ABA, which is recognized by the PYR/PYL/RCAR (PYRABACTIN RESISTANCE/PYRABACTIN RESISTANCE-LIKE/REGULATORY COMPONENT OF ABA RECEPTORS) intracellular soluble receptors. Hormone-receptor binding facilitates the formation of a protein complex with PP2C (PROTEIN PHOSPHATASE 2C) phosphatases, preventing the latter from inhibiting core ABA signaling protein kinases named SnRK2 [SUCROSE NON-FERMENTING 1 (SNF1)-RELATED PROTEIN KINASE 2]. In the absence of PP2Cs, SnRK2s activate themselves by autophosphorylation and can then phosphorylate specific transcription factors (TFs), such as bZIPs (BASIC LEUCINE ZIPPER), to induce the expression of stress-responsive genes (reviewed in Finkelstein, 2013; Sah et al., 2016).

Phosphorylation-triggered protein degradation is an important strategy common to animal and plant systems (reviewed in Filipčík et al., 2017; Vu et al., 2018; Bhaskara et al., 2019). In plants, it is crucial not only for the regulation of light signaling (Al-Sady et al., 2006; Shen et al., 2007; Yue et al., 2016) but also of several important players in the ABA pathway (Liu and Stone, 2010; Li et al., 2016; Chen et al., 2018; Li et al., 2018; Mizoi et al., 2019). The ubiquitin-proteasome system is a highly regulated mechanism to control protein levels in eukaryotic cells that targets for degradation proteins covalently linked to the polypeptide ubiquitin. For a protein to be recognized by the 26S proteasome, a chain of at least four ubiquitin monomers is needed, which is delivered to the substrate by an ATP-dependent enzymatic E1-E2-E3 conjugation cascade (reviewed in Sadanandom et al., 2012). E3 ubiquitin ligases are the most numerous and diverse of the three groups of enzymes, being responsible for substrate recognition and specificity. Substrate phosphorylation is one of the signals recognized by some types of E3 ubiquitin ligase complexes (reviewed in Deshaies, 1999).

The most studied A*rabidopsis thaliana* SR-related protein is SR45, a *bona fide* splicing factor (Ali et al., 2007) that plays an established role in ABA responses (Carvalho et al., 2010; Xing et al., 2015; Carvalho et al., 2016; Albaqami et al., 2019). Having been classified as a canonical SR protein for many years, SR45 is currently considered an SR-like protein (Barta et al. 2010) due to its two RS domains that flank a single RRM (Golovkin and Reddy, 1999). The only *SR45* loss-of-function mutant described to date, *sr45-1*, exhibits a variety of developmental phenotypes, including reduced plant size, late and bushy inflorescences, delayed root growth, as well as abnormal leaf and flower morphology (Ali et al., 2007). Moreover, *sr45-1* mutant seedlings are hypersensitive to ABA and glucose (Carvalho et al., 2010; Carvalho et al., 2016) as well as to high salinity (Albaqami et al., 2019). Seven different splice variants of the *A. thaliana SR45* gene are annotated, but only two have been characterized: *SR45*.*1* and *SR45*.*2*, which despite exhibiting very similar expression patterns, fulfill different functions. While both variants are able rescue the mutant’s glucose hypersensitivity (Carvalho et al., 2010), *SR45*.*2* rescues the root growth defect and *SR45*.*1* rescues the flower (Zhang and Mount, 2009), salt (Albaqami et al., 2019) and (partially) ABA (Xing et al., 2015) phenotypes.

Interestingly, the 4262 RNAs reported by Xing et al (2015) to bind SR45 include transcripts encoding 30% of all ABA signaling genes (Hauser et al., 2011). Furthermore, in addition to interacting with other splicing factors, including U1-70K (Golovkin and Reddy, 1999), U2AF (Day et al., 2012), three different U5 snRNP components and several SR proteins (Golovkin and Reddy, 1999; Tanabe et al., 2009; Zhang et al., 2014), SR45 has been found to interact with two proteins involved in ABA responses, the SNW/SKI-INTERACTING PROTEIN (SKIP) (Wang et al., 2012), a putative transcription factor conferring plant salt tolerance (Feng et al., 2015), and the SUA SUPPRESSOR OF ABI3-5 (SUA) protein (Mukhtar et al., 2011), which controls alternative splicing of *ABA-INSENSITIVE 3* (*ABI3*) (Sugliani et al., 2010).

Several reports of phosphorylation at specific SR45 residues in different plant tissues and stress conditions have been published (De La Fuente Van Bentem et al., 2006; De La Fuente Van Bentem et al., 2008; Umezawa et al., 2013; Wang et al., 2013; Zhang et al., 2014), with an early study showing that a LAMMER-type protein kinase, AFC2, can interact with and phosphorylate SR45 *in vitro* (Golovkin and Reddy, 1999). Moreover, Zhang et al. (2014) reported the relevance of phosphorylation of a single SR45 amino acid residue, T218, in both the regulation of flower development and alternative splicing of a direct mRNA target. On the other hand, a phosphoproteomics study found evidence of ABA-induced dephosphorylation of SR45 at a different residue, T264 (Wang et al., 2013). Given that SR45 transcript and splicing levels are reportedly unchanged by ABA treatment (Palusa et al., 2007; Cruz et al., 2014), these data strongly suggest that SR45 is regulated by ABA at the posttranslational level.

How SR45 is posttranslationally regulated by ABA and related stresses remains largely unknown. Here we report that the *A. thaliana* SR45 protein accumulates and is dephosphorylated at several residues in response to ABA. We find that ABA-mediated dephosphorylation of the protein reduces its ubiquitination and targeting for proteasomal degradation, thus leading to SR45 stabilization under ABA conditions. Our results also demonstrate that the T264 SR45 residue is sufficient to influence phosphorylation-dependent proteasomal degradation of the protein and thereby its function as a negative regulator of the ABA pathway.

## RESULTS

### Overexpression of the SR45 protein causes plant ABA hyposensitivity

We previously showed that loss of function of the *Arabidopsis SR45* gene leads to enhanced sensitivity to the ABA hormone at the cotyledon greening stage (Carvalho et al., 2010), with Xing et al. (2015) later reporting that overexpression of the *SR45*.*1* splice variant partially rescues this ABA greening inhibition phenotype. Given that the *SR45* gene produces at least two splice isoforms with distinct functions (Zhang and Mount, 2009; Albaqami et al., 2019), we decided to investigate the effect of expressing the genomic *SR45* fragment on the ABA response of transgenic seedlings during early development. To this end, we cloned the gene’s genomic fragment upstream of the eGFP sequence driven by either the endogenous *SR45* (pSR45::gSR45-eGFP) or the strong, constitutive *UBQ10* (pUBQ10::gSR45-eGFP) promoter (Supplemental Figure 1). These two constructs were independently transformed into *sr45-* mutant plants (Ali et al., 2007), with two complementation (C1 and C2) and two overexpression (OX1 and OX2) transgenic lines being isolated for the pSR45::gSR45-eGFP and pUBQ10::gSR45-eGFP constructs, respectively.

As seen in Figure 1A, RT-qPCR analysis using primers specific for the *SR45* gene (see Supplemental Figure 1) revealed that the OX1 and OX2 lines express about 15- and 30-fold higher transcript levels than the Col-0 wild type, respectively, while in the C2 line expression is enhanced by only ∼2.5 fold and C1 expresses similar levels to the wild type. As expected, transcript levels were barely detectable in the *sr45-1* knockout mutant. Similar results were obtained when primers specific for the *GFP* coding sequence were used (see Supplemental Figure 1) and the expression levels of the transgene were normalized to those of the C2 complementation line (Figure 1A). Western blot analysis using anti-GFP antibodies showed that the relative differences in *SR45-GFP* transcript among the transgenic lines were matched at the protein level, as both overexpression lines showed highly elevated amounts of SR45-GFP when compared to the complementation lines, with C2 exhibiting slightly higher levels of the fusion protein than C1 (Figure 1B).

**Figure 1.**
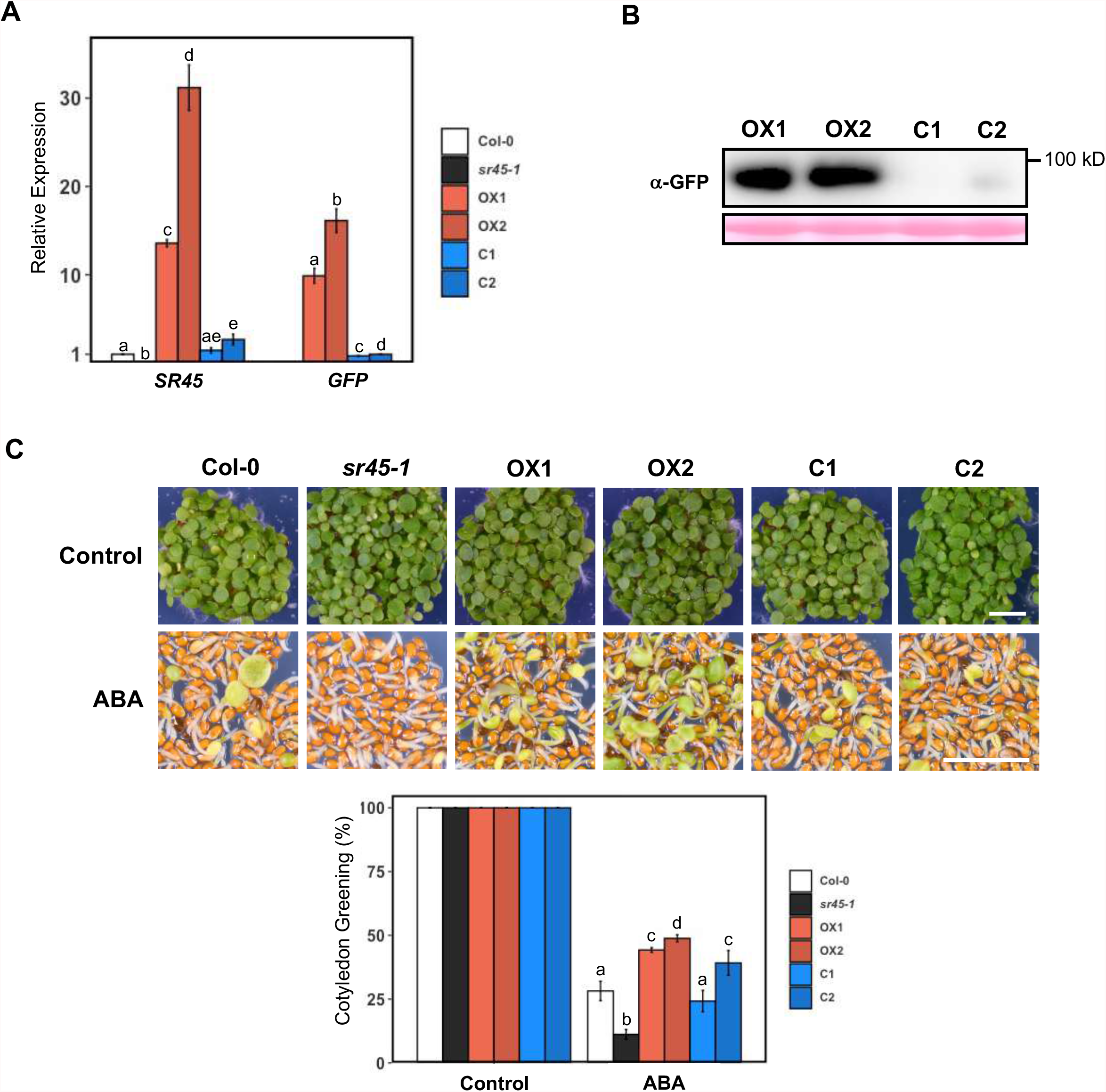
Effect of *SR45* overexpression on ABA signaling during cotyledon development. **(A)**RT-qPCR analysis of *SR45-GFP* transcript levels in 7-day-old pUBQ10::SR45-GFP/*sr45-1* (OX1 and OX2) and pSR45::SR45-GFP/*sr45-1* (C1 and C2) transgenic seedlings grown under control conditions, using *PEX4* as a reference gene and primers annealing to either the *SR45* or *GFP* sequences (see supplemental Figure 1). Transcript levels of either the Col-0 (*SR45* primers) or the C2 transgenic line (*GFP* primers) were set to 1. Results represent means ± SE (*n* = 3), and different letters indicate statistically significant differences between genotypes for each set of primers (P < 0.05; Student’s *t*-test). **(B)**Protein gel blot analysis using α-GFP antibodies of the SR45-GFP fusion protein in 7-day-old seedlings of overexpression (OX1 and OX2) and complementation (C1 and C2) transgenic lines grown under control conditions. A total of 30 ng of protein was loaded per sample, and Ponceau staining is shown as a loading control. Results are representative of at least 3 independent experiments. **(C)** Representative images and quantification of cotyledon greening in 7-day-old seedlings of the Col-0 wild type, the *sr45-1* mutant, the OX1 and OX2 overexpression lines and the C1 and C2 complementation lines grown under control conditions or in the presence of 0.5 μM ABA (means ± SE, *n* = 3). Different letters indicate statistically significant differences between genotypes under each condition (P < 0.05; Student’s *t*-test). Scale bar = 2.5 mm.

Phenotypical characterization of the transgenic lines revealed no defects in cotyledon development under control conditions and full rescue of the ABA hypersensitivity of the *sr45-1* mutant (Figure 1C). Moreover, all plant lines expressing significantly enhanced *SR45* mRNA levels (OX1, OX2 and C2) displayed reduced sensitivity to exogenous ABA. Notably, OX2 was the transgenic line that displayed stronger ABA hyposensitivity, correlating with higher expression of the *SR45-GFP* transgene. Our results show that SR45’s function as a negative regulator of ABA signaling during early seedling development is dependent on SR45 protein levels.

### ABA enhances SR45 protein levels

Previous work indicates that the expression levels and splicing pattern of the *SR45* gene are unchanged by ABA (Palusa et al., 2007; Cruz et al., 2014), but it remains unknown whether SR45 protein levels are regulated by the phytohormone. To investigate this, we treated seedlings from a transgenic complementation line with 2 µM ABA and followed SR45 protein levels for 3 hours by western blotting. Results revealed an evident accumulation of the SR45-GFP fusion protein over time in the ABA-treated samples, with the highest levels being detected after 180 minutes, while control samples harvested at the same timepoints showed no increase in SR45 protein levels (Figure 2A). Although endogenous *SR45* transcript levels are not ABA regulated (Palusa et al., 2007; Cruz et al., 2014), we confirmed that our transgene was also not transcriptionally affected by ABA. As seen in Supplemental Figure 2A, RT-qPCR analysis of the *SR45-GFP* transcript in SR45-GFP/*sr45-1* seedlings revealed no changes in expression levels upon ABA treatment.

**Figure 2.**
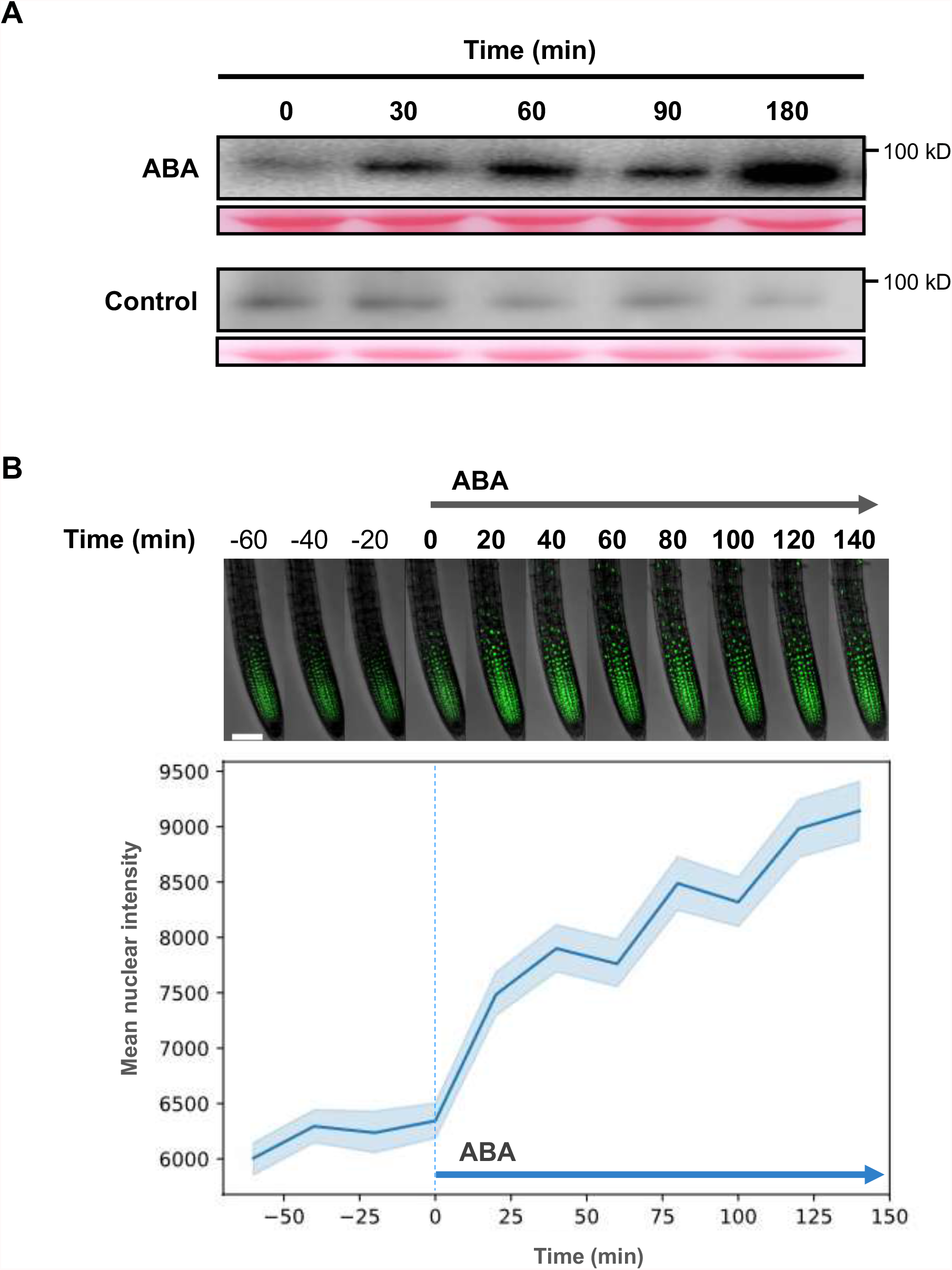
Effect of ABA on SR45 protein levels. **(A)**Protein gel blot analysis using α-GFP antibodies of the SR45-GFP fusion protein in 7-day-old seedlings of the C2 complementation transgenic line treated for 0, 30, 60, 90 or 180 minutes with 2 µM ABA. Control samples were treated with the equivalent volume of the solvent of the ABA solution (ethanol). A total of 40 ng of protein was loaded per sample, and Ponceau staining is shown as a loading control. Results are representative of at least 3 independent experiments. **(B)**Sum Z projection images (top) and fluorescence intensity quantification (bottom) of fast SR45-GFP accumulation in the primary root of 4-day-old seedlings of the C2 complementation transgenic line treated with 10 µM ABA observed by confocal microscopy. Scale bar = 100 μm. Line indicates mean values, with shaded region indicating the 95% confidence interval.

A similar trend of SR45-GFP accumulation was observed in transgenic seedlings treated with ABA when the fluorescence intensity was measured in primary roots by confocal microscopy. Figure 2B shows that addition of 10 µM ABA to the sample’s buffer increased the mean SR45-GFP fluorescence intensity per segmented nucleus by around 50% in 150 minutes, unequivocally demonstrating that the amounts of SR45 protein are upregulated by the ABA phytohormone.

### ABA dephosphorylates SR45 and enhances its levels in a SnRK2-dependent manner

In a phosphoproteomics study by Wang et al. (2013), the phosphorylation levels of SR45 were reported to decrease in response to an ABA treatment. To verify whether SR45 is dephosphorylated by ABA in our conditions, we treated seedlings from a complementation line with 2 µM ABA for 3 hours and compared the phosphorylation status of the SR45-GFP fusion protein with that from seedlings subjected to a mock treatment. As evident from the Phos-tag gel in Figure 3A, SR45 is markedly dephosphorylated in response to ABA. We next analyzed the phosphorylation levels of SR45 upon ABA treatment of transgenic plants expressing the pSR45::gSR45-eGFP construct in the *snrk2*.*2/3/6* triple mutant background. The SR45 dephosphorylation induced by ABA was severely impaired by loss of SnRK2 function, indicating that the switch in the protein’s phosphorylation status depends on ABA signaling (Figure 3A). These Phos-tag analyses revealed the occurrence of five different protein isoforms, while single bands had been detected in SDS-Page blots, thus showing that SR45 can be phosphorylated at multiple amino acid residues. We observed a clear SnRK2-dependent accumulation of the least phosphorylated form of the SR45 protein in response to ABA, with phospho-isoforms 3 and 4 accumulating in the ABA-treated *snrk2*.*2/3/6* line (Figure 3A). Importantly, when SR45 protein levels were analyzed in the same samples by western blotting, the SR45 protein accumulation observed upon ABA treatment (see also Figure 2A) was abolished in the *snrk2*.*2/3/6* mutant background (Figure 3B), indicating that the ABA-induced rise in SR45 levels fully depends on ABA signaling downstream of SnRK2s and suggesting a putative link between dephosphorylation of the SR45 protein and its accumulation. As in the complementation line (see Supplemental Figure 2A), RT-qPCR analysis of *SR45-GFP* transcript levels in SR45-GFP/*snrk2*.*2/3/6* seedlings revealed no changes upon ABA treatment (Supplemental Figure 2B).

**Figure 3.**
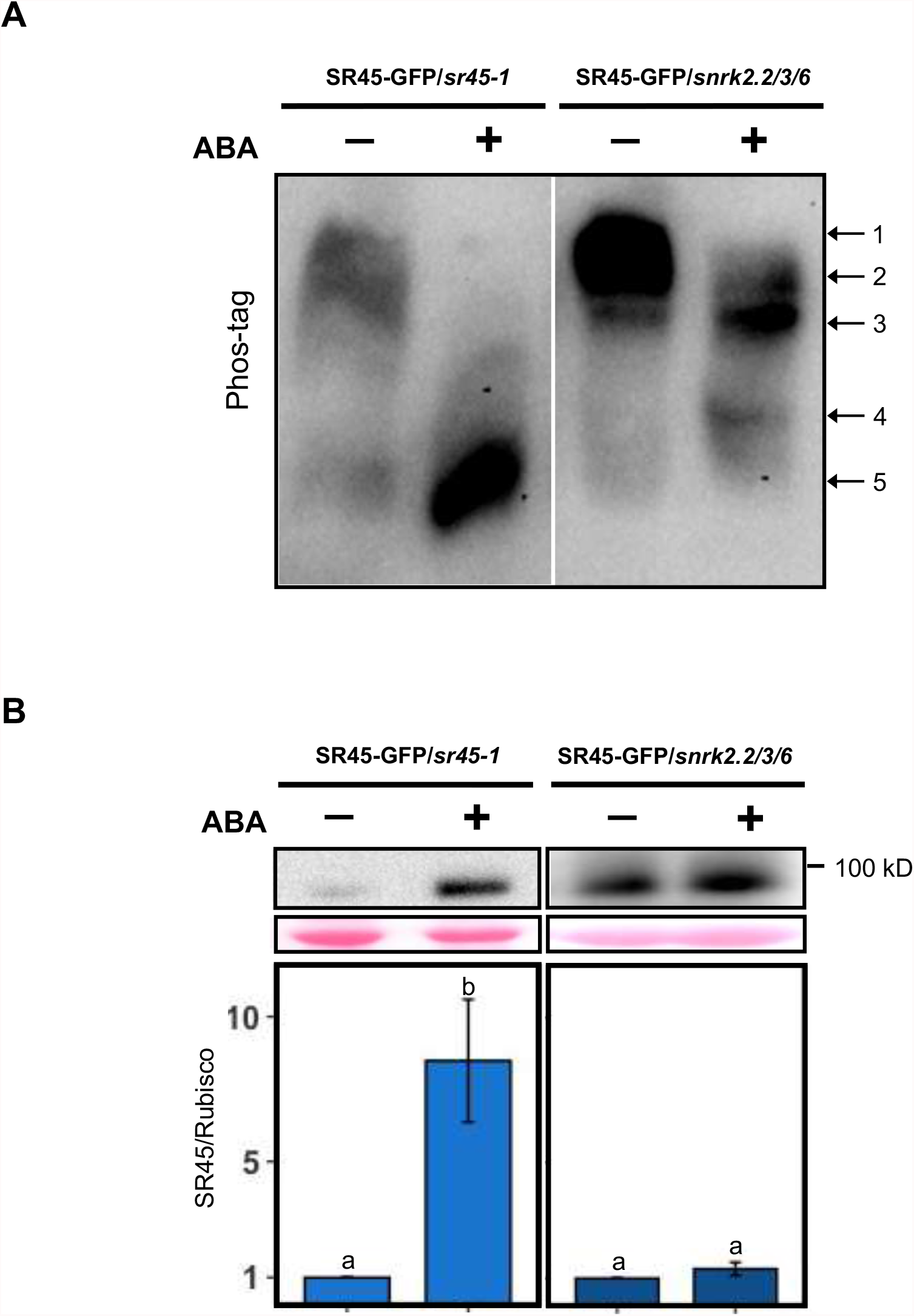
Effect of ABA and loss of SnRK2 function on SR45 protein phosphorylation and amounts. Phos-tag **(A)** and protein **(B)** gel blot analyses using α-GFP antibodies of the SR45-GFP fusion protein in 7-day-old seedlings of the C2 complementation (SR45-GFP/*sr45-1*) line and a transgenic line expressing the pSR45::gSR45-GFP construct in the *snrk2*.*2/3/6* mutant background (SR45-GFP/*snrk2*.*2/3/6*) treated for 180 minutes with 2 µM ABA. Control samples were treated with the equivalent volume of the solvent of the ABA solution (ethanol), and a total of 40 ng of protein were loaded per sample. In **(A)**, arrows indicate phosphorylated forms of SR45, and the results are representative of at least 3 independent experiments. In **(B)**, bands were quantified and relative protein levels determined using the Ponceau loading control as a reference, with results representing means ± SE (*n* = 4), control conditions set to 1, and different letters indicating statistically significant differences between treatments for each genotype (P < 0.05; Student’s *t*-test).

### ABA stabilizes SR45 by reducing protein ubiquitination and proteasomal degradation

Having established that SR45 is ABA regulated at the posttranslational level, we hypothesized that the different amounts of SR45 protein observed under control and ABA conditions were due to distinct stability of the protein. To test this, we treated seedlings with the potent proteasome inhibitor MG132 before exposure to ABA and determination of SR45-GFP protein levels by western blotting (Figure 4A). Notably, pre-treatment with MG132 resulted in a significant increase of SR45 levels only in the absence of ABA, suppressing the difference in SR45 content between the control and ABA conditions (Figure 4A). This indicates increased targeting of SR45 for proteasomal degradation under control conditions, strongly suggesting that the ABA-dependent SR45 protein accumulation is due to increased stability of the SR45 protein under ABA conditions.

**Figure 4.**
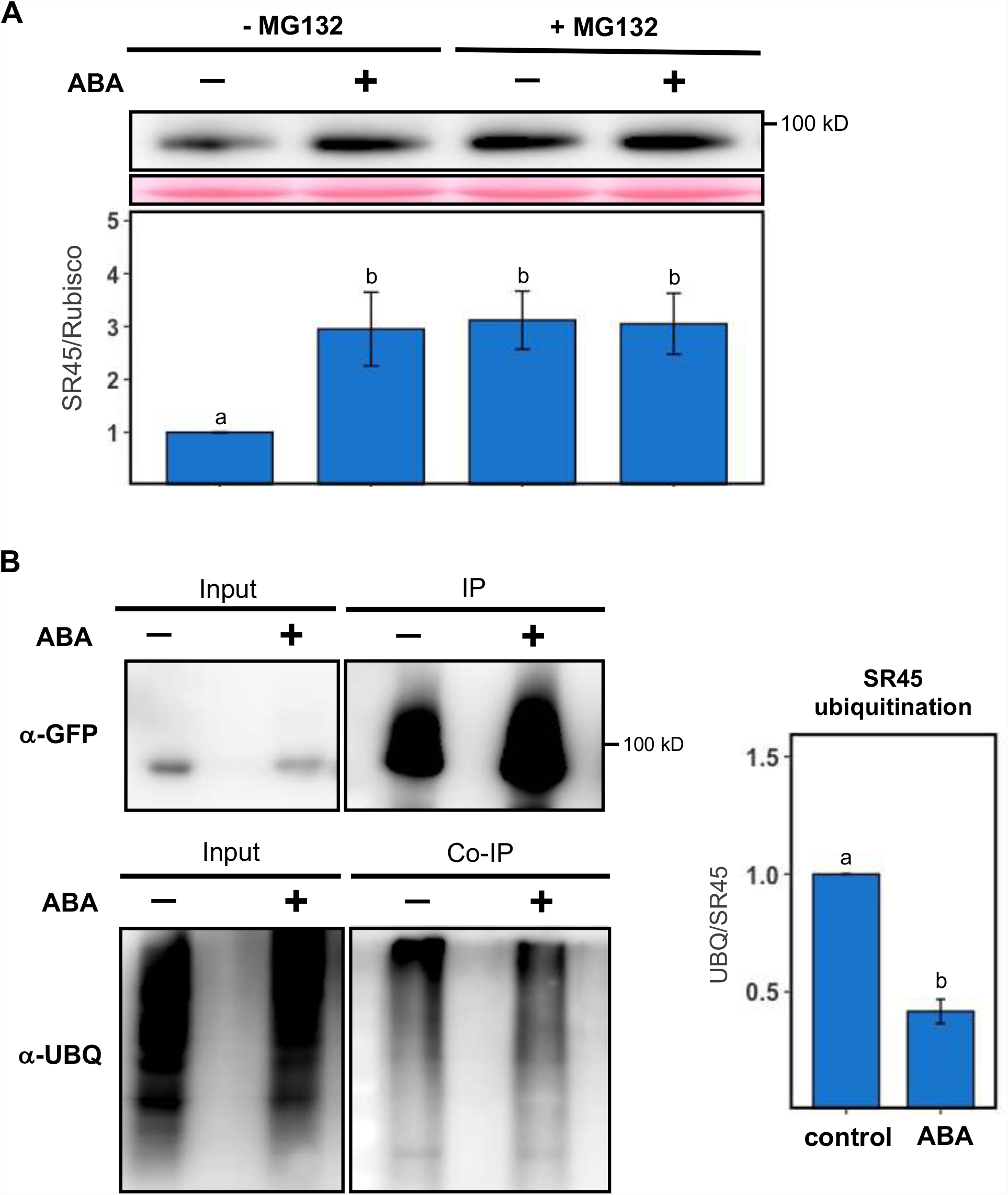
Effect of ABA on SR45 protein stability and ubiquitination. **(A)** Protein gel blot analysis using α-GFP antibodies of the SR45-GFP fusion protein in 7-day-old seedlings of the C2 complementation line pre-treated with MG132 and subjected to a 180-minute treatment with 2 µM ABA. Control samples (-MG132 or -ABA) were treated with the equivalent volume of the solvent of the MG132 or ABA solutions (DMSO or ethanol, respectively). A total of 40 ng of protein were loaded per sample. Bands were quantified and relative protein levels determined using the Ponceau loading control as a reference, with control conditions set to 1. Results represent means ± SE (*n* = 4), and different letters indicate statistically significant differences between treatments (P < 0.05; Student’s *t*-test). **(B)** Protein gel blot analysis of the SR45-GFP fusion protein immunoprecipitated from extracts of 7-day-old seedlings of the C2 complementation line treated for 180 minutes with 2 µM ABA using α-GFP (IP) or α-UBQ11 (Co-IP) antibodies. Control samples were treated with the equivalent volume of the solvent of the ABA solution (ethanol). Equal volumes of both the input fraction (Input) and the IP were loaded. Signals were quantified and the UBQ/SR45-GFP ratio determined, with control conditions set to 1. Results represent means ± SE (*n* = 3), and different letters indicate statistically significant differences between treatments (P < 0.05; Student’s *t*-test).

To verify that the differences in SR45 stability correlate with different ubiquitination levels of the protein, we immunoprecipitated SR45-GFP (IP) from protein extracts of seedlings treated with ABA and checked for the presence of ubiquitin conjugates (Figure 4B). As expected, for the same amount of IP loaded in the SDS-Page gel, more immunoprecipitated SR45-GFP was retrieved in ABA-treated samples when compared to the control, but a higher accumulation of ubiquitin was detected in control compared to ABA pull-downs. Quantification of the ubiquitin signal and normalization to the amounts of immunoprecipitated SR45-GFP showed that the ubiquitination levels of the immunoprecipitated SR45-GFP are reduced by more than half under ABA conditions (Figure 4B). Immunoprecipitation of control 35S::GFP transgenic seedlings showed residual ubiquitin conjugates bound to GFP alone (Supplemental Figure 3). Together, these results show a higher degree of both SR45 ubiquitination and degradation by the ubiquitin-proteasome system under control conditions, with the protein being less ubiquitinated and more stable upon exposure to ABA.

### Phosphorylation of T264 residue controls SR45 protein ubiquitination and degradation

Given that phosphorylation-dependent protein ubiquitination and degradation is a known mechanism to rapidly control protein levels in both animal and plant systems (reviewed in Filipčík et al., 2017; Vu et al., 2018; Bhaskara et al., 2019), we next asked whether SR45 stability is dependent on the phosphorylation status of the protein. To address this question, we generated transgenic *Arabidopsis* lines expressing the pUBQ10::gSR45-eGFP construct mutated at a threonine (T) phosphoresidue reported by Wang et al. (2013). We changed this T264 residue either to an alanine (A), an amino acid that cannot be phosphorylated (phosphomutant lines), or to aspartic acid (D), which is structurally similar to a phosphorylated threonine and thus mimics constitutive phosphorylation (phosphomimetic lines).

To investigate whether SR45 phosphorylation at the T264 residue affects ubiquitination of the protein, we immunoprecipitated SR45-GFP from protein extracts of overexpression (OX1), phosphomutant (PMut1) and phosphomimetic (PMim1) transgenic seedlings grown in control conditions and checked for the presence of ubiquitin in the IPs (Figure 5A). Quantification of the respective signals and calculation of the ubiquitin/SR45 ratio showed much higher ubiquitination levels in the pulldowns from the phosphomimetic line when compared with the overexpression and phosphomutant lines, with SR45-GFP immunoprecipitated from the latter line, in which T264 is never phosphorylated, showing the lowest degree of ubiquitination (Figure 5A). These results indicated that phosphorylation of the T264 residue promotes SR45 ubiquitination.

**Figure 5.**
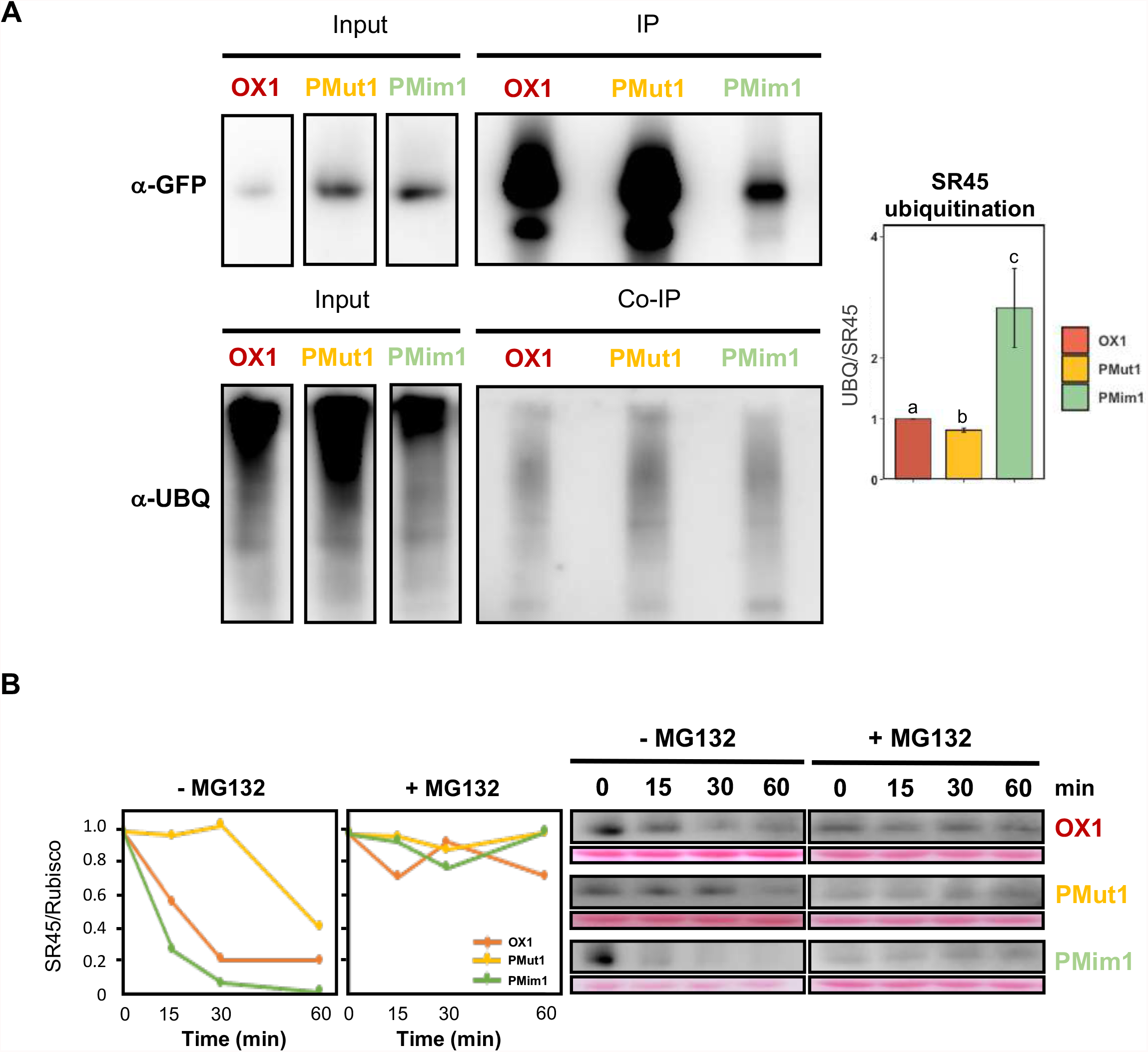
Effect of T264 phosphorylation on SR45 ubiquitination and degradation. **(A)** Protein gel blot analysis of the SR45-GFP fusion protein immunoprecipitated from extracts of 7-day-old seedlings of the OX1 overexpression (pUBQ10::SR45-GFP/*sr45-1*), PMut1 phosphomutant (pUBQ10::SR45-GFP_T264A/*sr45-1*) and PMim1 phosphomimetic (pUBQ10::SR45-GFP_T264D/*sr45-1*) transgenic lines grown in control conditions using α-GFP (IP) or α-UBQ11 (Co-IP) antibodies. Equal volumes of both the input fraction (Input) and the IP were loaded. Signals were quantified and the UBQ/SR45-GFP ratio determined, with control conditions set to 1. Results represent means ± SE (*n* = 3), and different letters indicate statistically significant differences between genotypes (P < 0.05; Student’s *t*-test). **(B)** Protein gel blot analysis using α-GFP antibodies of the SR45-GFP fusion protein in 7-day-old seedlings of the OX1 overexpression, PMut1 phosphomutant and PMim1 phosphomimetic transgenic lines supplemented or not with MG132 and left at room temperature for 0, 15, 30 or 60 minutes. Control samples (-MG132) were treated with the equivalent volume of the solvent of the MG132 solution (DMSO), and a total of 20 ng of protein were loaded per sample. Bands were quantified and relative protein levels determined using the Ponceau loading control as a reference, with time 0 set to 1. Results are representative of at least 3 independent experiments.

To verify that T264 phosphorylation also results in rapid destabilization of SR45, we next followed degradation of the protein along time. As observed in Figure 5B, while in the control overexpression line SR45 decayed to about 60% of its initial levels in 15 minutes, the protein was noticeably more rapidly degraded in the phosphomimetic line, falling to nearly 20% in the same timeframe. By contrast, SR45 was markedly more stable in phosphomutant extracts, beginning to decay only after 30 minutes and reducing its amounts to only about half after an hour. Addition of the MG132 inhibitor to the protein extracts showed that the observed decay of the SR45 protein in all plant lines is proteasome dependent (Figure 5C). We thus concluded that T264 phosphorylation controls SR45 protein degradation via the ubiquitin-proteasome system — dephosphorylation of the protein at this residue results in reduced levels of ubiquitination and therefore enhanced protein stability.

### Phosphorylation of T264 residue controls ABA-induced SR45 protein accumulation and plant ABA sensitivity

Finally, we analyzed the effect of SR45 phosphorylation at the T264 residue on ABA responses. First, to test ABA-driven SR45 protein accumulation, we treated seedlings from the OX1 overexpression, the PMut1 phosphomutant and the PMim1 phosphomimetic transgenic lines with 2 µM ABA and assessed SR45-GFP levels after 3 hours by western blotting. As shown in Figure 6A, the accumulation of SR45-GFP under ABA conditions observed in the C2 complementation line (see Figure 2 and Figure 3B) was reproduced with OX1, an overexpression line in which the transgene is driven by a different promoter, reinforcing posttranslational regulation of SR45 by ABA. By contrast, the ABA-induced SR45 protein accumulation was abolished in both the transgenic line where SR45 cannot undergo phosphorylation at T264 (PMut1) and in the line where SR45 is constitutively phosphorylated at this residue (PMim1). We actually observed a slight decrease in the SR45-GFP protein levels upon ABA treatment in the phosphomutant and phosphomimetic lines (Figure 6A) consistent with reduced transcript levels (Supplemental Figure 4), possibly due to ABA downregulation of *UBQ10* promoter activity. This transcriptional downregulation of expression of the transgene by ABA was also observed in an independent set of phospho-transgenic lines, OX2, PMut2 and PMim2 (Supplemental Figure 5A), where again a rise in SR45 protein levels in response to ABA treatment was observed only in the overexpression line (Supplemental Figure 5B). These results demonstrate that a switch in the phosphorylation status is required for SR45 protein accumulation in response to ABA exposure, corroborating the notion that the ABA-induced increase in the SR45 protein depends on its dephosphorylation by the hormone.

**Figure 6.**
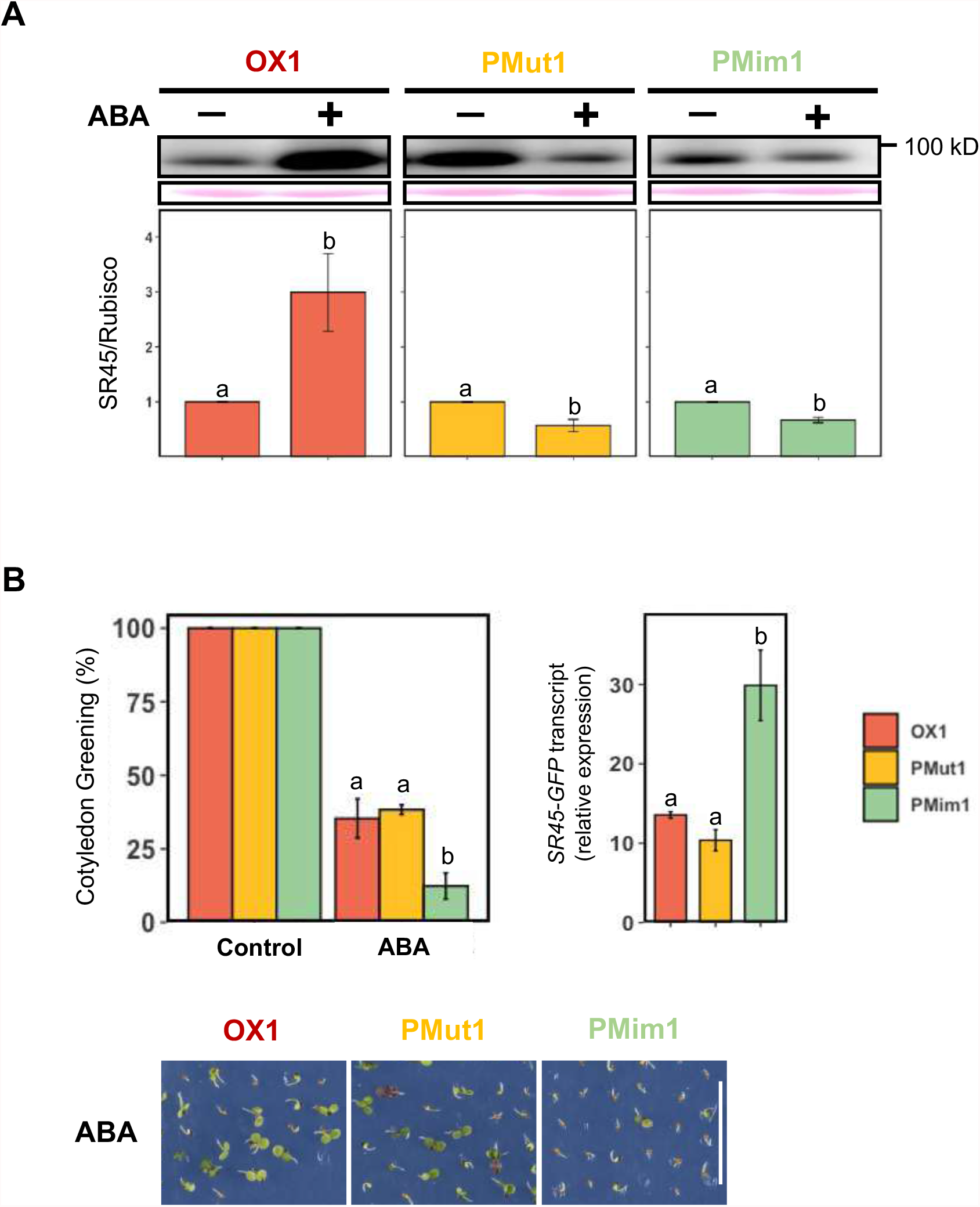
Effect of T264 phosphorylation on ABA-dependent SR45 protein accumulation and cotyledon development. **(A)** Protein gel blot analysis using α-GFP antibodies of the SR45-GFP fusion protein in 7-day-old seedlings of the OX1 overexpression (pUBQ10::SR45-GFP/*sr45-1*), PMut1 phosphomutant (pUBQ10::SR45-GFP_T264A/*sr45-1*) and PMim1 phosphomimetic (pUBQ10::SR45-GFP_T264D/*sr45-1*) transgenic lines treated for 180 minutes with 2 µM ABA. Control samples were treated with the equivalent volume of the solvent of the ABA solution (ethanol), and a total of 20 ng of protein were loaded per sample. Bands were quantified and relative protein levels determined using the Ponceau loading control as a reference, with results representing means ± SE (*n* = 3), control conditions set to 1, and different letters indicating statistically significant differences between treatments for each genotype (P < 0.05; Student’s *t*-test). **(B)** Cotyledon greening percentages of 7-day-old seedlings of the OX1 overexpression, PMut1 phosphomutant and PMim1 phosphomimetic transgenic lines grown under control conditions or in the presence of 0.5 μM ABA, with representative images of ABA conditions (scale bar = 1 cm), and RT-qPCR analysis of *SR45-GFP* transcript levels in the same seedlings (control conditions), using *PEX4* as a reference gene and primers annealing to the *GFP* sequence (see Supplemental Figure 1). Results represent means ± SE (*n* = 3), and different letters indicate statistically significant differences between genotypes (P > 0.05; Student’s *t*-test).

Having gained molecular insight into the effects of the T264 mutation on SR45, we then asked whether it would impact the plant’s physiological response to ABA and affect sensitivity to the hormone during cotyledon greening. Figure 6B shows that phenotypical characterization of the transgenic OX1, PMut1 and PMim1 lines under control conditions revealed no differences in cotyledon development. However, while the OX1 and Pmut1 lines displayed a similar reduction in cotyledon greening when grown in ABA, the PMim1 phosphomimetic seedlings showed a higher sensitivity to ABA, despite the fact that this particular line expressed about twice the levels of the SR45-GFP mRNA (Figure 6B). A similar result was obtained when independent transgenic lines (OX2, PMut2 and PMim2) were isolated and analyzed, with the PMim2 phosphomimetic line being clearly the most sensitive to ABA (Supplemental Figure 6). In this case, OX2 was less sensitive to ABA when compared to PMut2, most likely because of markedly higher *SR45-GFP* transcript levels in the former transgenic line (Supplemental Figure 6). In conclusion, the fact that SR45 phosphomimetic transgenic lines, in which the T264 residue is constitutively phosphorylated, exhibit enhanced ABA sensitivity indicates that SR45 depends on dephosphorylation to enhance its levels and exert its role as a negative regulator of ABA responses during early plant development.

## DISCUSSION

We show here that overexpression of the *Arabidopsis SR45* gene causes reduced plant sensitivity to the phytohormone ABA. While Carvalho et al. (2010) observed complete rescue of the partially ABA-dependent *sr45-1* glucose phenotype upon overexpression of either the *SR45*.*1* or *SR45*.*2* splice variants, Xing et al. (2015) found that overexpressing the *SR45*.*1* splice variant partially reverts *sr45-1* ABA hypersensitivity during early seedling development. Neither study had identified an ABA hyposensitive phenotype upon *SR45* overexpression, perhaps due to the fact that individual *SR45* splice variants were tested independently instead of the genomic fragment. Seven *SR45* transcripts are currently annotated and at least *SR45*.*1* and *SR45*.*2* are known to fulfill different biological roles (Zhang and Mount, 2009; Albaqami et al., 2019), rendering it likely that a combination of SR45 splice forms is required for the protein’s role as a negative regulator of the ABA pathway. Alternatively, the ability to reduce ABA sensitivity could arise from the use of different promoters to drive the *SR45* transgene. In our overexpression lines, the genomic construct is driven by the *UBQ10* promoter, while previous work used *35S*, whose activity can vary in different organs and under abiotic stress (Kiselev et al., 2021). Nevertheless, we recently also found opposite ABA and ABA-related phenotypes for the loss-of-function mutant and *Arabidopsis* lines expressing the SCL30a SR protein under the control of the 35S promoter (Laloum et al., 2021).

SR proteins are established phosphoproteins, and several phosphorylation residues have been annotated for SR45. Moreover, a phosphoproteomics study listed SR45 as displaying reduced phosphorylation under ABA conditions (Wang et al. 2013). In agreement, we found that upon ABA exposure SR45 is dephosphorylated at several residues in a SnRK2-dependent manner, indicating that ABA signaling downstream of SnRK2 kinases is either activating phosphatases that dephosphorylate SR45 or deactivating kinases that phosphorylate SR45 under control conditions. Interestingly, we detected ABA-induced accumulation of a specific SR45 phospho-isoform (isoform 4 in Figure 3A) in the *snrk2*.*2/3/6* background, suggesting that this SR45 (de)phosphorylation event is occurring either due to ABA signaling upstream of SnRK2s or independently of these core ABA pathway components. SnRK2-independent ABA signaling, in which PP2C phosphatases dephosphorylate other kinases activating different branches of ABA signaling, has been previously reported (Brandt et al., 2012). Our findings fit with the general model in which dephosphorylation negatively regulates ABA signaling (reviewed in Yang et al., 2017). In fact, several PP2C phosphatases are known to negatively regulate ABA signaling, with the corresponding loss-of-function mutants displaying ABA hypersensitive phenotypes (Merlot et al., 2001; Yoshida et al., 2006; Nishimura et al., 2007), as observed for SR45.

Importantly, we found that the SR45 protein accumulates upon ABA treatment in a SnRK2-dependent manner and that the phosphorylation status of the protein controls this ABA-induced increase in SR45 levels. Indeed, using phosphomutant and phosphomimetic lines in which the SR45 protein is never phosphorylated or constitutive phosphorylation is mimicked at T264, a previously reported ABA-specific SR45 dephosphorylation residue (Wang et al., 2013), we show that: *i*) a switch in the SR45 phosphorylation status is required for ABA induction of SR45 protein accumulation, *ii*) dephosphorylation of the T264 residue upon ABA exposure results in both reduced ubiquitination levels and lower proteasomal degradation rates of SR45, and *iii*) SR45 phosphorylation at T264 enhances plant ABA sensitivity at the early seedling stage.

Our results demonstrate functional and physiological relevance for ABA-mediated posttranslational modification of an SR-related splicing factor that negatively regulates ABA responses during early plant development (Carvalho et al., 2010; Xing et al., 2015). We show that SR45 is phosphorylated under control conditions, with phosphorylation of a single residue being able to modulate the protein’s ubiquitination levels and the amounts of SR45 that are targeted to degradation by the ubiquitin-proteasome system. Upon exposure to ABA, downstream signaling triggers SR45 dephosphorylation, thus reducing both the ubiquitination and degradation rates of the protein. Consistent with ABA stabilization of the SR45 protein, we also demonstrate that the protein’s role as negative regulator of the ABA pathway is dependent on SR45 protein levels, pointing to ABA-induced SR45 accumulation as a mechanism of negative autoregulation of ABA signaling. Also in agreement with our model (Figure 7), SR45 phosphomimetic lines are more sensitive to ABA — when T264 dephosphorylation is prevented, SR45 is degraded at higher rates resulting in lower amounts of SR45 protein and enhanced seedling sensitivity to ABA; given that SR45 is dephosphorylated by ABA, both the overexpression and the phosphomutant lines display a decreased ABA response.

**Figure 7.**
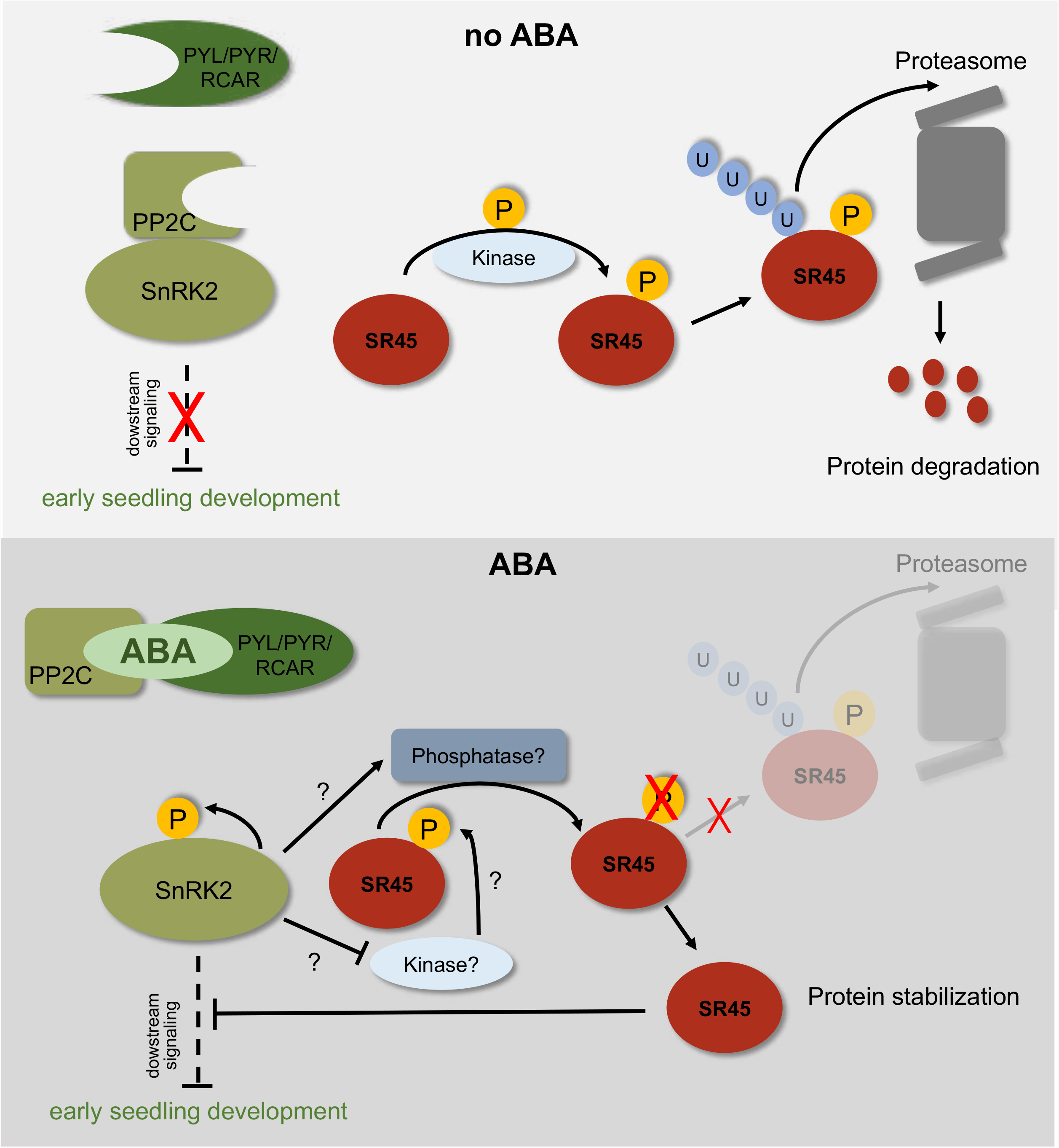
Model of ABA-mediated SR45 regulation of early seeding development. In the absence of ABA, PP2Cs inhibit SnRK2 activity, thus blocking ABA signaling and its inhibition of early seedling development. Under these conditions, SR45 is phosphorylated by (an) unknown kinase(s), triggering SR45 ubiquitination and proteasomal degradation. When ABA accumulates in the cell, the hormone binds to the PYL/PYR/RCAR receptors creating a complex with PP2C, thus derepressing SnRK2s that are then able to activate themselves through autophosphorylation and induce downstream signaling. ABA signaling either activates (a) phosphatase(s) or inactivates (a) kinase(s) that dephosphorylate SR45, leading to its deubiquitination and stabilization. The increase in SR45 protein levels then results in negative regulation of ABA signaling, alleviating its inhibition of early seedling development.

In plants, protein phosphodegradation is crucial in the regulation of the phytochrome interacting factor 5 (PIF5) and PIF3 transcription factors involved in light signaling (Al-Sady et al., 2006; Shen et al., 2007; Yue et al., 2016), but has also been shown to regulate directly key ABA signaling components, including the *Arabidopsis* PYR/PYLs receptors (Chen et al., 2018), DEHYDRATION-RESPONSIVE ELEMENT-BINDING PROTEIN 2A (DREB2A) (Mizoi et al., 2019) and GUANINE NUCLEOTIDE EXCHANGE FACTOR 1 (RopGEF1) (Li et al., 2016; Li et al., 2018). Moreover, the RING-type E3 ligase KEEP ON GOING (KEG) regulates itself via ABA-dependent phosphodegradation, thereby stabilizing the ABA-INSENSITIVE 5 (ABI5) transcription factor, which is ubiquitinated by this E3 ligase (Liu and Stone, 2010). As for SR proteins, although their phosphorylation-dependent degradation has not yet been reported, a couple of studies in humans have looked into both phosphorylation and proteasomal degradation of these RNA-binding proteins. Breig and Baklouti (2013) found that mutating an AKT signaling phosphorylation site has no effect on proteasomal degradation of the human SR protein SRSF5, while acetylation downregulates phosphorylation levels and promotes degradation of SRSF2, suggesting a link between acetylation and phosphorylation in regulating the levels of this SR protein (Edmond et al., 2011).

In line with our findings, POLYUBIQUITIN 3 (UBQ3) is a reported interactor of the *Arabidopsis* SR45 (Kim et al., 2013). SR45 has also been reported to interact with the products of the At2g43770 and At1g10580 genes (Zhang et al., 2014), which are predicted to contain WD40 domains (Zhang et al., 2008). WD40 proteins function as potential substrate receptors of CUL4 E3 ubiquitin ligases (Lee et al., 2008), and it is interesting to note that two components of these E3 ligases are also reported negative regulators of ABA signaling (Lee et al., 2010). However, phosphorylated substrates are usually recognized by F-box proteins, which are part of the the Skp1-Cullin-F-box (SCF) complex, a different major type of E3 ubiquitin ligases (reviewed in Deshaies, 1999; Sadanandom et al., 2012). Further biochemical and genetic studies should allow the identification of both the kinases or phosphatases and the E3 ligase complexes regulating SR45 protein levels and unveil the precise molecular mechanisms leading to SR45 dephosphorylation and accumulation in response to ABA.

## METHODS

### Plant Materials

The *Arabidopsis thaliana* Colombia ecotype (Col-0) was used as the wild type in all experiments. The *sr45-1* knockout homozygous line was originally isolated in the Duque lab (Carvalho et al. 2010). The *snrk2*.*2/3/6* triple mutant was kindly provided by P.L. Rodríguez (Universidad Politécnica de Valencia, Spain).

### Generation of transgenic lines

To generate the pSR45::gSR45-eGFP constructs, a fragment including the *SR45* promoter and its genomic sequence was isolated by PCR using primers annealing 1252 bp upstream and 2756 bp downstream of the *SR45* start codon (Supplemental Table 1) and Col-0 DNA as a template. The *SR45* amplicon was subcloned into pGEM®-T Easy vector and incorporated in a GFP-tagged version of the binary pBA002 vector using the BspEI/PacI restriction sites. The construct was then used to transform the *Arabidopsis sr45-1* and *snrk2*.*2/3/6* mutants by the floral dip method (Clough and Bent, 1998) using *Agrobacterium tumefaciens* strain GV3101. The full-length *SR45* fragment (excluding the stop codon) was amplified using primers with attB1/attB2 MultiSite Gateway sites (Supplemental Table 1). An entry clone was generated introducing the amplicon into a pDONR™221 (Invitrogen) by recombination. The entry clone *SR45* in pDONR™221 along with the vectors with the ubiquitin-10 promoter (UBQ10) in pDONR™P4-P1R (Invitrogen) and eGFP in pDONR™P2R-P3 (Invitrogen) was recombined with the destination vector pHm43GW to generate the pUBQ10::gSR45-eGFP overexpression construct according to Hartley et al. (2000). To generate SR45 variants carrying the T264A and T264D mutations, site-directed mutagenesis was performed using *Pfu* DNA polymerase on *SR45* in the pDONR™221 clone using different primer pairs (Supplemental Table 1). The resulting clones were further recombined with *UBQ10* in pDONR™P4-P1R and *eGFP* in pDONR™P2R-P3 in the pHm43GW destination vector as described above for the overexpression construct. Each construct was independently transformed into *sr45-1* mutant plants by agroinfiltration. All experiments with transgenic lines were conducted on isolated T3 homozygous lines.

### Plant growth

For all phenotypical assays, seeds were surface sterilized for 10 minutes with 50 % (v/v) bleach and 0.02 % (v/v) Tween X-100 under continuous shaking and then washed three times in sterile water. Approximately 100 seeds per genotype were plated in triplicate on half-strength MS medium (0.5X MS basal salt mix Duchefa Biochemie, 0.5 mM *myo*-inositol, 2.5 mM MES, and 0.8 % agar, adjusted to pH 5.7) supplemented or not with 0.5 μM ABA (S-ABA Duchefa Biochemie). The plates were wrapped in aluminum foil and stored at 4 °C for 3 days. After stratification, plates were transferred to a growth chamber set to continuous light conditions (100 μmol m ^−2^ s ^−1^) and 22 °C. The percentage of green seedlings was calculated over the total number of germinated seeds (displaying an emerged radicle) after 7 days. The average of the percentages was calculated per genotype and statistical differences between the genotypes were assessed using a Student’s *t*-test.

For protein extraction, seedlings were grown for 5 days in 0.5X MS agar medium under continuous light conditions (100 μmol m^−2^ s^−1^) at 22 °C. The seedlings were then transferred to liquid 0.5X MS medium and grown with constant shaking for 48 hours in the same growth conditions. At this stage, control (ethanol) or 2 μM ABA (S-ABA Duchefa Biochemie; ethanol stock) treatments were applied for specific amounts of time (0, 30, 60, 90 or 180 minutes). To inhibit proteasomal degradation, 50 μM MG132 (Sigma, C2211; DMSO stock) or DMSO (control) were added 60 minutes prior to the onset of the ABA treatment. The plant material was harvested and frozen at -80 °C for protein extraction.

To compare the RNA levels of all transgenic lines, plants were also grown for 5 days in 0.5X MS agar medium under continuous light and transferred to liquid 0.5X MS medium for 48 hours with constant shaking. To test for transcriptional ABA regulation, 2-day-old seedlings were grown on a paper filter and placed on 0.5X MS agar medium plates under continuous light conditions. The seedlings were then transferred to 0.5X MS agar medium supplemented with 1 μM ABA or ethanol (control) and grown for 180 minutes before harvesting.

### RNA Extraction and RT-qPCR Analyses

Total RNA was extracted from *Arabidopsis* seedlings using the innuPREP Plant RNA kit (Analytik Jena BioSolutions). All RNA samples were treated with DNAseI (Promega) and cDNA was synthesized using the SuperScript™ III Reverse Transcriptase (Invitrogen) and oligo (dT)_18_ primers. RT-qPCR was performed using an ABI QuantStudio-384 instrument (Applied Biosystems) and the Luminaris Color HiGreen qPCR Master Mix, high ROX (Thermo Scientific) on 2.5 μL of cDNA (diluted 1:20) per 10-μL reaction volume, containing 300 nM of each specific primer (Supplemental Table 1). Cycle threshold (Ct) values were adjusted for each gene in each biological replicate. Relative expression values were generated for the target gene using a log2 transformation, normalizing the gene Ct values to the housekeeping control gene *PEX4* (*PEROXIN4)*, according to Vandesompele et al. (2002). The average value of the relative expression of each gene in all the biological replicates was calculated per sample. Statistical differences between the average relative expressions of each sample were inferred using a Student’s *t*-test.

### Microscopic Analyses

For confocal microscopy analysis, *sr45-1* seedlings expressing the pSR45::gSR45-eGFP construct were mounted on slides in a vacuum grease/coverslip reservoir as described in Rizza et al. (2019) in 0.25X MS. Seedlings were imaged for an hour of pretreatment, and then a buffer exchange was performed to 0.25X MS + 10 µM ABA and imaging was resumed. Confocal images were acquired on an upright Leica SP8 microscope using a HC PL APO CS2 20x/0.75 DRY objective. The emission laser and detection windows were as follows: GFP - 488nm laser, 493nm - 568nmdetection; Gain: 200. A custom Fiji (Schneider et al., 2012; Rueden et al., 2017) plugin, ‘Simple_auto_segmentation.py’, was developed using CLIJ2 (Haase et al., 2020) to perform segmentation and fluorescence quantification per nucleus. Source code and installation instructions are available at https://github.com/JimageJ/ImageJ-Tools. This preprocesses images for segmentation with difference of Gaussian and tophat filters followed by thresholding using Otsu’s method. A watershed and connected components analysis are used to split and identify objects in the threshold image. Any objects lost in watershed are added back in and a non-zero dilation and multiplication is used to expand the object map into the original binary map to produce the final segmentation.

### Protein Extraction, Western Blot and Phos-tag Assays

Frozen 7-day-old *Arabidopsis* seedlings subjected to prior treatments were ground with a mortar and pestle under liquid nitrogen. Total protein was extracted in an extraction buffer containing 50 mM Tris-HCl, 150 mM NaCl, 1 mM EDTA, 0.5% Triton™ X-100(Sigma-Aldrich), 1 tablet of Complete Protease Inhibitor Cocktail (Roche) and 1 tablet of PHOSSTOP (Roche). The extract was centrifuged at 18,000 *g* for 10 minutes at 4 °C and the protein content of the supernatant determined using the Bradford method (Bio-Rad) in a spectrophotometer measuring the absorbance at 595 nm and then eluted in Laemmli 2X buffer with 8 % β-mercaptoethanol (95 °C for 5 minutes). After determining the protein concentration in each sample, equal amounts of protein were resolved in 8 % SDS/polyacrylamide gels. The proteins were then transferred to PVDF membranes (Immobilon-P; Millipore) and subsequently blocked with 5 % nonfat dry milk for 2 hours. The membranes were probed overnight at 4 °C with anti-GFP primary antibodies (Roche 11814460001; 1:1000) and then with anti-mouse peroxidase-conjugated secondary antibodies (Jackson Immunoresearch # 115-035-146; diluted 1:4000) for 2 hours at room temperature. All antibodies were diluted in TBS buffer (25 mM Tris-HCl pH 7.4, and 137 mM NaCl) supplemented with 1 % nonfat dry milk. After incubating with the antibodies, the membranes were washed with TBS containing 0.05 % Tween^®^ 20 (Sigma-Aldrich) for 40 minutes. The peroxidase activity associated with the membrane was visualized by enhanced chemiluminescence. The intensity of the protein bands was quantified using ImageJ software, normalizing protein levels to the Rubisco large subunit visualized in membranes stained with Ponceau (0.1 % Ponceau S Sigma-Aldrich in 5 % acetic acid). The statistical differences between the average protein levels of each sample were inferred using a Student’s *t*-test.

To separate the different SR45 phospho-isoforms under control and ABA conditions, Mn^2+^-Phos-tag™ SDS-PAGE was performed. An equivalent amount of total protein was loaded and resolved in a 6 % acrylamide gel with 50 µM MnCl_2_ and 25 µM pf Phos-tag™ (Wako Pure Chemicals Industries). Each sample was supplemented with 1:10 MnCl_2_ to minimize the “smiling” effect before the run. The gel was washed twice for 20 minutes with transfer buffer (25 mM Tris, 192 mM glycine, 0.1 %, SDS and 20 % ethanol) supplemented with 1mM EDTA, followed by an additional wash of regular transfer buffer for 10 minutes. The proteins were wet-transferred to a PVDF membrane (Immobilon-P; Millipore) for 2.5 hours at 100 V. The membrane was blocked and probed with antisera as mentioned above.

### SR45-GFP Immunoprecipitation and Protein Degradation Assay

Total protein was extracted from approximately 100 mg of 7-day-old *Arabidopsis* seedlings with 800 μL of the extraction buffer described above. The extract was centrifuged at 18,000 *g* for 10 minutes at 4 °C and the supernatant incubated for 1 hour at 4 °C with continuous agitation with 100 μL of Sepharose beads (Sigma-Aldrich). The extract was then further centrifuged at 500 *g* for 2 minutes, and the input fraction was removed and eluted in Laemmli 2x buffer for 5 minutes at 95 °C. After incubation with 20 μL of GFP-Trap^®^ Agarose beads (ChromoTek) for 1.5 h, the beads were washed 3 times with cold extraction buffer for 10 minutes each wash and the proteins eluted from the beads in 30 μL of Laemmli 2x buffer for 5 minutes at 95 °C. Finally, 15 μL of the immunoprecipitate and 20 μL of the input fraction were loaded in separate SDS-Page gels, with blotting performed as described above. For detection of UBQ conjugates, the membranes were stripped for 15 minutes with Western BLoT Stripping Buffer (Takara) following the manufacturer’s instructions and reprobed with Ubiquitin11 (Agrisera AS08307; 1/10000) and anti-rabbit peroxidase-conjugated secondary antibodies (Amersham Pharmacia; diluted 1/10000). The intensity of the protein bands was quantified using ImageJ software. Statistical differences between the average ratios of each sample were inferred using a Student’s *t*-test.

The degradation rate of the SR45 protein was assessed in protein extracts from 7-day-old seedlings grown in control conditions supplemented with 50 μM MG132 or DMSO (control) and left to degrade at room temperature for 0, 15, 30 and 60 minutes. The protein extracts were loaded in an SDS-Page gel and blotting performed as described above.

## SUPPLEMENTAL INFORMATION

**Supplemental Figure 1. Constructs for generation of the overexpression and complementation transgenic lines**.

Schematic representation of the pUBQ10::gSR45-eGFP and pSR45::gSR45-eGFP constructs. The *UBQ10* and the *SR45* promoters are shown in orange and blue, respectively, and the Egfp sequence in green. Exons are shown in black, the 5’ UTR in white, and introns are represented by black lines. The cloning scar sequences are shown immediately downstream of the promoter and/or of the last exon. The location of the primers used to detect the transgene is indicated by the arrow pairs.

**Supplemental Figure 2. Effect of ABA and loss of SnRK2 function on *SR45-GFP* transcript levels**.

**(A)** RT-qPCR analysis of *SR45-GFP* transcript levels in 2-day-old seedlings of the C2 complementation (SR45-GFP/*sr45-1*) line **(A)** or of a transgenic line expressing the pSR45::gSR45-GFP construct in the *snrk2*.*2/3/6* mutant background (SR45-GFP/*snrk2*.*2/3/6*) **(B)** treated for 180 minutes with 1 µM ABA, using *PEX4* as a reference gene and primers annealing to the *GFP* sequence (see Supplemental Figure 1). Control samples (set to 1) were treated with the equivalent volume of the solvent of the ABA solution (ethanol). Results represent means ± SE (*n* = 3), with no statistically significant differences being found between treatments for each set of primers (P > 0.05; Student’s *t*-test).

**Supplemental Figure 3. Ubiquitination levels of the GFP protein**.

Protein gel blot analysis of the GFP protein immunoprecipitated from extracts of 7-day-old seedlings of a 35S::GFP transgenic line treated for 180 minutes with 2 µM ABA using α-GFP (IP) or α-UBQ11 (Co-IP) antibodies. Control samples were treated with the equivalent volume of the solvent of the ABA solution (ethanol). Equal volumes of both the input fraction (Input) and the IP were loaded.

**Supplemental Figure 4. Effect of ABA on *SR45-GFP* transcript levels in the OX1, PMut1 and PMim1 transgenic lines**.

RT-qPCR analysis of *SR45-GFP* transcript levels in 2-day-old seedlings of the OX1 overexpression (pUBQ10::SR45-GFP/*sr45-1*), PMut1 phosphomutant (pUBQ10::SR45-GFP_T264A/*sr45-1*) and PMim1 phosphomimetic (pUBQ10::SR45-GFP_T264D/*sr45-1*) transgenic lines treated for 180 minutes with 1 µM ABA, using *PEX4* as a reference gene and primers annealing to the *GFP* sequence (see Supplemental Figure 1). Control samples (set to 1) were treated with the equivalent volume of the solvent of the ABA solution (ethanol). Results represent means ± SE (*n* = 3), with different letters indicating statistically significant differences between treatments for each genotype (P > 0.05; Student’s *t*-test).

**Supplemental Figure 5. Effect of ABA on *SR45-GFP* transcript and protein levels in the OX2, PMut2 and PMim2 transgenic lines**.

**(A)**RT-qPCR analysis of *SR45-GFP* transcript levels in 2-day-old seedlings of the OX2 overexpression (pUBQ10::SR45-GFP/*sr45-1*), PMut2 phosphomutant (pUBQ10::SR45-GFP_T264A/*sr45-1*) and PMim2 phosphomimetic (pUBQ10::SR45-GFP_T264D/*sr45-1*) transgenic lines treated for 180 minutes with 1 µM ABA, using *PEX4* as a reference gene and primers annealing to the *GFP* sequence (see Supplemental Figure 1). Control samples (set to 1) were treated with the equivalent volume of the solvent of the ABA solution (ethanol). Results represent means ± SE (*n* = 3), with different letters indicating statistically significant differences between treatments for each genotype (P > 0.05; Student’s *t*-test). **(B)**Protein gel blot analysis using α-GFP antibodies of the SR45-GFP fusion protein in 7-day-old overexpression (OX2), phosphomutant (PMut2) and phosphomimetic (PMim2) transgenic seedlings treated for 180 minutes with 2 µM ABA. Control samples were treated with the equivalent volume of the solvent of the ABA solution (ethanol), and a total of 20 ng of protein were loaded per sample. Bands were quantified and relative protein levels determined using the Ponceau loading control as a reference, with results representing means ± SE (*n* = 3), control conditions set to 1, and different letters indicating statistically significant differences between treatments for each genotype (P < 0.05; Student’s *t*-test).

**Supplemental Figure 6. Physiological phenotypes of the OX2, PMut2 and PMim2 transgenic lines**.

Cotyledon greening percentages of 7-day-old seedlings of the OX2 overexpression, PMut2 phosphomutant and PMim2 phosphomimetic transgenic lines grown under control conditions or in the presence of 0.5 μM ABA, with representative images of ABA conditions (scale bar = 1 cm), and RT-qPCR analysis of *SR45-GFP* transcript levels in the same seedlings (control conditions), using *PEX4* as a reference gene and primers annealing to the *GFP* sequence (see Supplemental Figure 1). Results represent means ± SE (*n* = 3), and different letters indicate statistically significant differences between genotypes (P > 0.05; Student’s *t*-test).

**Supplemental Table 1. Sequences of the Primers Used in this study**

## AUTHOR CONTRIBUTIONS

R.A.-M., A.M.J. and P.D. designed the research, R.A.-M., D.S. and J.R. performed the experiments, and A.M.J. and P.D. supervised the work. R.A.-M. and P.D. wrote the manuscript and prepared the figures and tables. All authors contributed to the interpretation of results, critically reviewed the manuscript and approved its final version.

## ACKNOWLEDGMENTS

We thank P.L. Rodríguez for the *snrk2*.*2/3/6* triple mutant, L. Margalha for expert biochemistry advice, and V. Nunes for excellent plant care at the Instituto Gulbenkian de Ciência (IGC) Plant Facility.

