## Supplemental Information for "ABA signaling prevents phosphodegradation of the *Arabidopsis* SR45 splicing factor to negatively autoregulate inhibition of early seedling development"

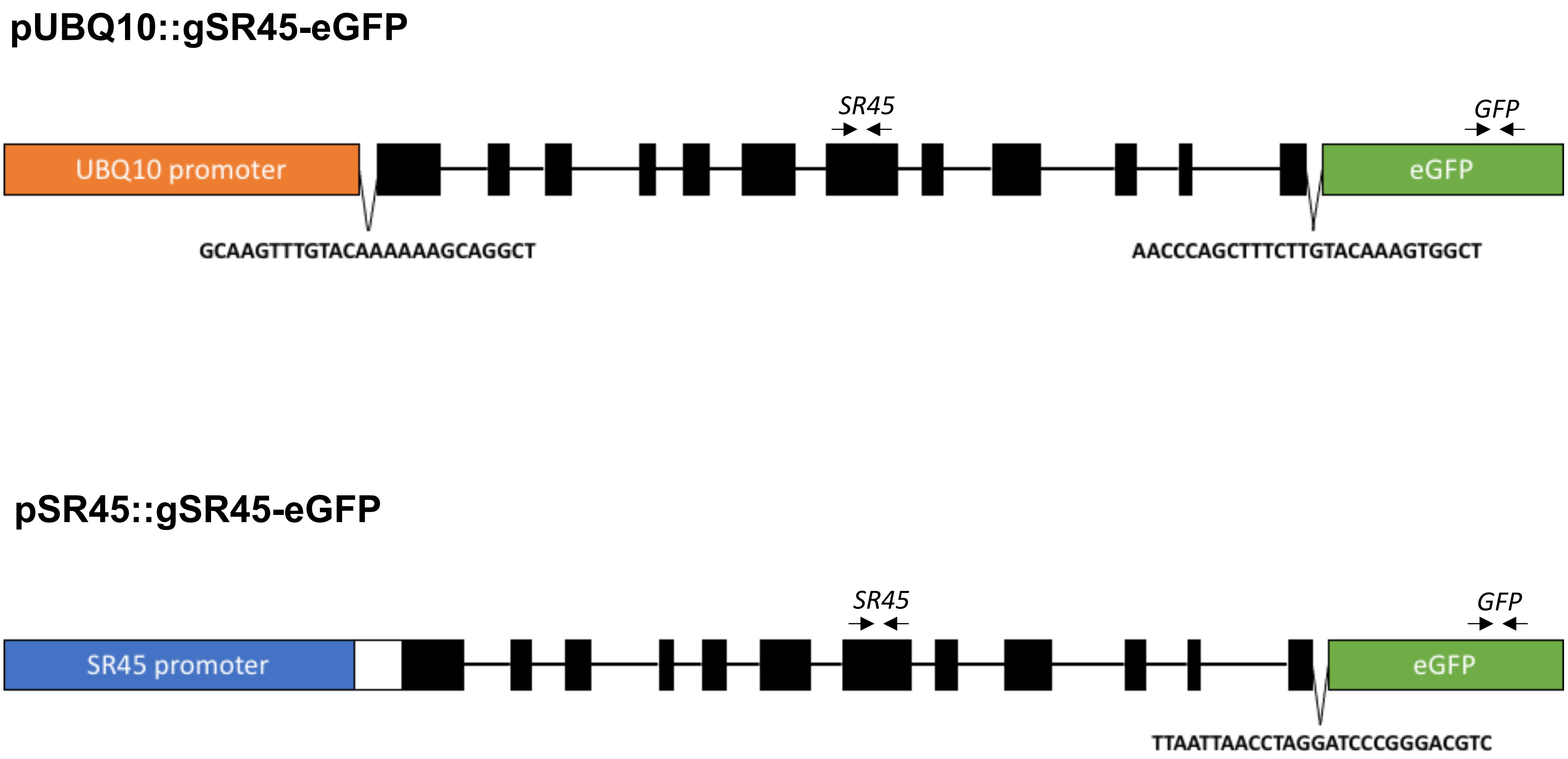


**Supplemental Figure 1. Constructs for generation of the overexpression and complementation transgenic lines.**

Schematic representation of the pUBQ10::gSR45-eGFP and pSR45::gSR45-eGFP constructs. The *UBQ10* and the *SR45* promoters are shown in orange and blue, respectively, and the eGFP sequence in green. Exons are shown in black, the 5' UTR in white, and introns are represented by black lines. The cloning scar sequences are shown immediately downstream of the promoter and/or of the last exon. The location of the primers used to detect the transgene is indicated by the arrow pairs.

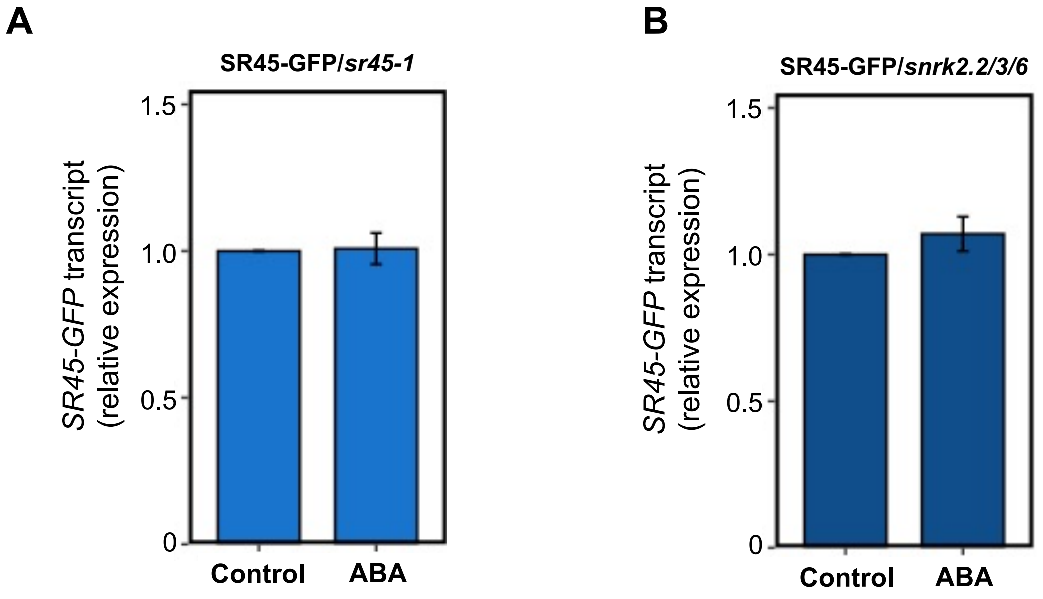


**Supplemental Figure 2. Effect of ABA and loss of SnRK2 function on *SR45-GFP* transcript levels.**

RT-qPCR analysis of *SR45-GFP* transcript levels in 2-day-old seedlings of the C2 complementation (SR45-GFP/*sr45-1*) line **(A)** or of a transgenic line expressing the pSR45::gSR45-GFP construct in the *snrk2.2/3/6* mutant background (SR45-GFP/*snrk2.2/3/6*) **(B)** treated for 180 minutes with 1 µM ABA, using *PEX4* as a reference gene and primers annealing to the *GFP* sequence (see Supplemental Figure 1). Control samples (set to 1) were treated with the equivalent volume of the solvent of the ABA solution (ethanol). Results represent means ± SE ($n$ = 3), with no statistically significant differences being found between treatments for each set of primers ($P > 0.05$; Student's *t*-test).

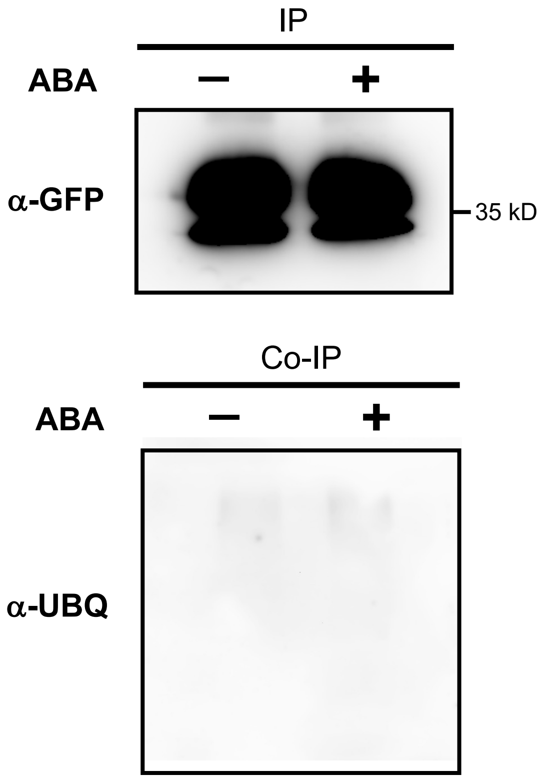


**Supplemental Figure 3. Ubiquitination levels of the GFP protein.**

Protein gel blot analysis of the GFP protein immunoprecipitated from extracts of 7-day-old seedlings of a 35S::GFP transgenic line treated for 180 minutes with 2 μM ABA using α-GFP (IP) or α-UBQ11 (Co-IP) antibodies. Control samples were treated with the equivalent volume of the solvent of the ABA solution (ethanol). Equal volumes of both the input fraction (Input) and the IP were loaded.

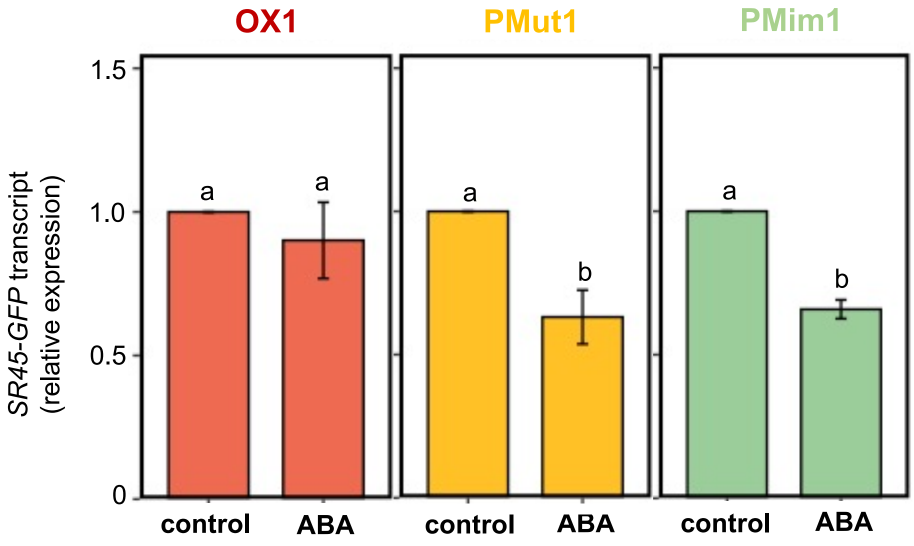


**Supplemental Figure 4. Effect of ABA on *SR45-GFP* transcript levels in the OX1, PMut1 and PMim1 transgenic lines.**

RT-qPCR analysis of *SR45-GFP* transcript levels in 2-day-old seedlings of the OX1 overexpression (pUBQ10::SR45-GFP/*sr45-1*), PMut1 phosphomutant (pUBQ10::SR45-GFP_T264A/*sr45-1*) and PMim1 phosphomimetic (pUBQ10::SR45-GFP_T264D/*sr45-1*) transgenic lines treated for 180 minutes with 1 μM ABA, using *PEX4* as a reference gene and primers annealing to the *GFP* sequence (see Supplemental Figure 1). Control samples (set to 1) were treated with the equivalent volume of the solvent of the ABA solution (ethanol). Results represent means ± SE ($n$ = 3), with different letters indicating statistically significant differences between treatments for each genotype ($P > 0.05$; Student's $t$-test).

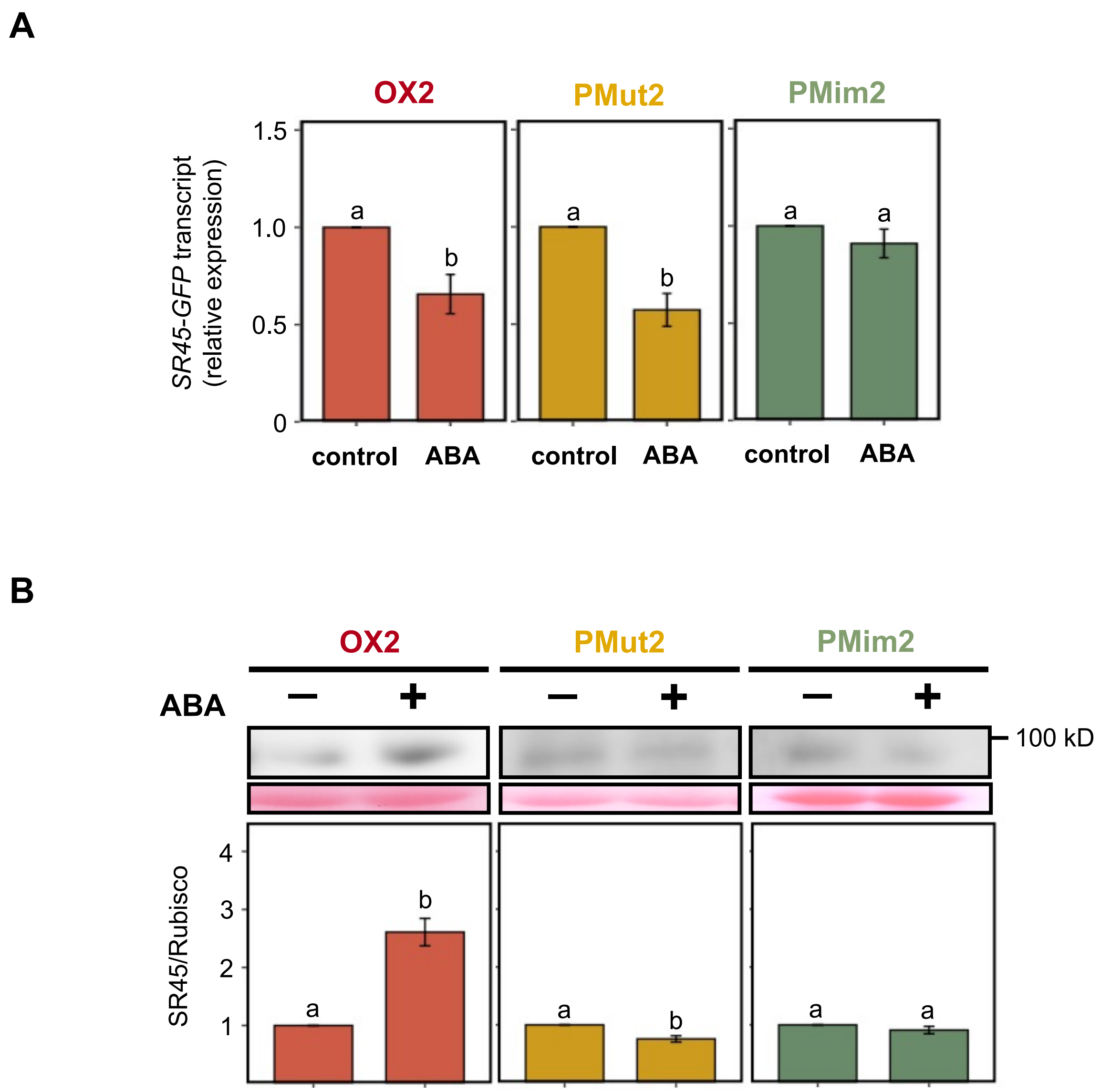


**Supplemental Figure 5. Effect of ABA on *SR45-GFP* transcript and protein levels in the OX2, PMut2 and PMim2 transgenic lines.**

**(A)** RT-qPCR analysis of *SR45-GFP* transcript levels in 2-day-old seedlings of the OX2 overexpression (pUBQ10::SR45-GFP/*sr45-1*), PMut2 phosphomutant (pUBQ10::SR45-GFP_T264A/*sr45-1*) and PMim2 phosphomimetic (pUBQ10::SR45-GFP_T264D/*sr45-1*) transgenic lines treated for 180 minutes with 1 μM ABA, using *PEX4* as a reference gene and primers annealing to the *GFP* sequence (see Supplemental Figure 1). Control samples (set to 1) were treated with the equivalent volume of the solvent of the ABA solution (ethanol). Results represent means ± SE ($n$ = 3), with different letters indicating statistically significant differences between treatments for each genotype ($P > 0.05$; Student's *t*-test).

**(B)** Protein gel blot analysis using $\alpha$-GFP antibodies of the SR45-GFP fusion protein in 7-day-old overexpression (OX2), phosphomutant (PMut2) and phosphomimetic (PMim2) transgenic seedlings treated for 180 minutes with 2 μM ABA. Control samples were treated with the equivalent volume of the solvent of the ABA solution (ethanol), and a total of 20 ng of protein were loaded per sample. Bands were quantified and relative protein levels determined using the Ponceau loading control as a reference, with results representing means ± SE ($n$ = 3), control conditions set to 1, and different letters indicating statistically significant differences between treatments for each genotype ($P < 0.05$; Student's *t*-test).

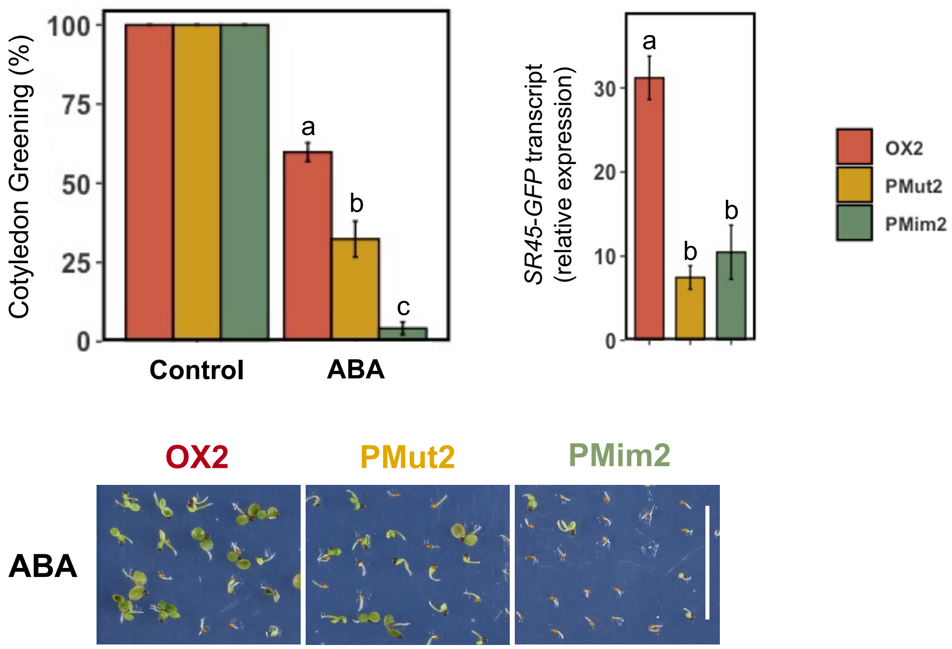


**Supplemental Figure 6. Physiological phenotypes of the OX2, PMut2 and PMim2 transgenic lines.**

Cotyledon greening percentages of 7-day-old seedlings of the OX2 overexpression, PMut2 phosphomutant and PMim2 phosphomimetic transgenic lines grown under control conditions or in the presence of 0.5 μM ABA, with representative images of ABA conditions (scale bar = 1 cm), and RT-qPCR analysis of *SR45-GFP* transcript levels in the same seedlings (control conditions), using *PEX4* as a reference gene and primers annealing to the *GFP* sequence (see Supplemental Figure 1). Results represent means ± SE ($n$ = 3), and different letters indicate statistically significant differences between genotypes (P > 0.05; Student's $t$-test).

### Supplemental Table 1. Primers used in this study.

| GENE | PRIMER | SEQUENCE |
|---|---|---|
| **RT-qPCR analyses** | | |
| *SR45* | SR45qF1 | 5'-GCGATCACCTGATTCTCCC-3' |
| | SR45qR1 | 5'-AGATCTATATCGTCTTGGAGG-3' |
| *GFP* | GFPqF1 | 5'-CCACTACCAGCAGAACACCC-3' |
| | GFPqR1 | 5'-GCTCAGGTAGTGGTTGTCGG-3' |
| *PEX4* | PEX4qF1 | 5'-TTACGAAGGCGGTGTTTTTC-3' |
| | PEX4qR1 | 5'-GGCGAGGCGTGTATACATTT-3' |
| **Generating T264A mutation in *gSR45* by site-directed mutagenesis** | | |
| | gSR45T264AF | 5'-TACGTCTAGGAGGAGCATCACCGCGACGGCG-3' |
| | gSR45T264AR | 5'-CGCCGTCGCGGTGATGCTCCTCCTAGACGTA-3' |
| **Generating T264D mutation in *gSR45* by site-directed mutagenesis** | | |
| | gSR45T264DF | 5’-CCTACGTCTAGGAGGATCATCACCGCGACGGCGGA-3’ |
| | gSR45T264DR | 5’-TCCGCCGTCGCGGTGATGATCCTCCTAGACGTAGG-3’ |
| **Cloning *pSR45:gSR45* into pBA002/eGFP vector** | | |
| | ProSR45F | 5’-AA*TCCGGA*CCCTGAAGAAGCAGTTTCATGAGG-3’ |
| | gSR45R | 5’-TT*TTAATTAA*AGTTTTACGAGGTGGAGGTGGTG-3’ |
| **Cloning *gSR45* into pDONR 221 vector** | | |
| | gSR45221F | 5’-*GGGGACAAGTTTGTACAAAAAAGCAGGCT*ATGGCGAAACCAAGTCGTGG-3’ |
| | gSR45221R | 5’-*GGGGACCACTTTGTACAAGAAAGCTGGGTT*AGTTTTACGAGGTGGAGGTG-3’ |

Restriction and recombination sites are shown in italics and underlined.
